# Active learning reveals underlying decision strategies

**DOI:** 10.1101/239558

**Authors:** Paula Parpart, Eric Schulz, Maarten Speekenbrink, Bradley C. Love

## Abstract

One key question is whether people rely on frugal heuristics or *full-information* strategies when making preference decisions. We propose a novel method, *model-based active learning*, to answer whether people conform more to a rank-based heuristic (Take-The-Best) or a weight-based full-information strategy (logistic regression). Our method eclipses traditional model comparison techniques by using information theory to characterize model predictions for how decision makers should actively sample information. These analyses capture how sampling affects learning and how learning affects decisions on subsequent trials. We develop and test model-based active learning algorithms for both Take-The-Best and logistic regression. Our findings reveal that people largely follow a weight-based learning strategy rather than a rank-based strategy, even in cases where their preference decisions are better predicted by the Take-The-Best heuristic. This finding suggests that people may have more refined knowledge than is revealed by their preference decisions, but which can be revealed by their information sampling behavior. We argue that model-based active learning is an effective and sensitive method for model selection that expands the basis for model comparison.

## Introduction

How do people decide between two alternatives? This question is as fundamental to studies of judgment and decision making as its answer is controversial (Todd & Gigerenzer, 2000). Whereas some researchers propose people only require few pieces of information for good decision making (Gigerenzer & Brighton, 2009; Marewski, Gaissmaier, & Gigerenzer, 2010; Şimşek, 2013), others have described them as integrating all available evidence (Arkes, Dawes, & Christensen, 1986). One of the core questions in this debate concerns the way in which people look up and integrate information (Bröder & Schiffer, 2003; Newell & Shanks, 2003).

Imagine you have to decide between two restaurants. Both restaurants differ on several binary features (for example, one is in walkable distance, the other is not). One decision strategy you could apply is to weigh the features by their importance and add them up; such as full-information strategies like regression or weighted additive models would do (Payne, Bettman, & Johnson, 1993, WADD). For each restaurant, this strategy would compute a weighted sum, and the restaurant with the larger sum is chosen. Alternatively, you might use a simpler strategy and base your decision on a single feature only. This is what the Take-The-Best heuristic (Gigerenzer & Goldstein, 1996, TTB) does: it ranks features by their validity (Martignon & Hoffrage, 1999), and chooses the restaurant that is preferred by the highest ranked feature. If that feature does not discriminate between restaurants, then the second feature is considered, and so forth^1^. Weighing and adding all features is a *compensatory* strategy, whereas TTB is a *non-compensatory strategy* (Martignon & Hoffrage, 1999). Compensatory strategies have the property that one feature’s implied decision can be compensated for by combinations of other features. Linear or logistic regression are typical examples of such strategies. In contrast, once a discriminating feature has been found, the non-compensatory TTB heuristic ignores all other features to make decisions. In this way, the most powerful discriminating feature *C_k_* outweighs any combination of the subsequent features *C_k+_*_1_*,…, C_k+n_*, for all *k* (Gigerenzer & Goldstein, 1999). In fact, it is assumed that a decision maker utilizing the TTB heuristic does not look up any features further down the rank order once a higher ranked feature points towards favoring one option - a property which has earned TTB the label “fast and frugal” (Gigerenzer, 2004). Hence, while a regression model would assume people are tuned to the feature weights, the TTB heuristic assumes that they rely on feature rank orders.

What strategy do people use? Many of the empirically derived arguments in favor of TTB are based on showing that TTB can outperform more complex, compensatory models at out-of-sample predictions (Czerlinski, Gigerenzer, & Goldstein, 1999; Gigerenzer & Brighton, 2009). Even though showing that heuristics can outperform more complex strategies is a necessary precursor for establishing their ecological validity, it is psychologically not the same as actually establishing that people apply these strategies. Furthermore, although the initial studies leading to the discussion of TTB’s ecological validity produced repeated evidence for “one good reason” decision strategies (Tversky, 1969), later studies have produced mixed results (Hilbig, 2010; Newell, Weston, & Shanks, 2003). The evidence for heuristic or full-information strategies is still inconclusive, and derives from a homogeneous set of model testing methods. Furthermore, these methods mostly rely on passive model testing, e.g., where stimuli are already pre-selected by the experimenter, not necessarily reflecting how learning and decision making takes place in the real world. We argue that novel model comparison approaches are needed - and that active information sampling may be a sensitive method to add to our model comparison toolkit. A related theoretical argument is that the common way of assessing people’s cognitive strategies in over-simplified tests may not always reveal what knowledge they have at their disposal.

We propose *model-based active learning* as a model discrimination method to assess cognitive model classes (Cohn, Ghahramani, & Jordan, 1996). The central idea is that if a cognitive agent prefers one decision strategy over another, this should also be reflected in the way she acquires information in a self-directed manner. If an agent has evolved to prefer a certain class of models as her means to learn a cognitive representation in a particular environment, then the way she sequentially selects information should (at least partially) reflect this representation. For example, if an agent has come to apply TTB, then —intuitively— she should try to gather information about feature rank orders. What most previous decision making studies have in common is that they study people’s decision making in static, passive and highly controlled experiments. Yet, in order to answer the crucial question about what information people hold in memory and how they look up knowledge when making decisions, we believe one has to also look at an earlier stage of the process -at the stage of learning the relevant information (see also Coenen, Nelson, & Gureckis, 2017). We argue that stronger evidence for people’s use of either TTB or a full-information-like strategies comes from the way they actively acquire information. Building on this intuition, we describe a general method to derive active learning versions of traditional models of learning and decision making, by concentrating on one-step-ahead uncertainty sampling (Cohn et al., 1996; Settles, 2010). Deriving active learning versions for both Take-The-Best and logistic regression, we put these two models to a test in a self-directed sampling task in which participants are able to select binary comparisons to learn about the underlying structure in an “Alien Olympics” game. Thus, we are able to observe both classic decision making behavior as well as active learning behavior. Our results show that logistic regression does not only predict participants’ choices in a binary prediction task better than TTB, but —more importantly— that self-directed learning behavior is also better described by a model-based active learning version of logistic regression than Take-The-Best. Participants seem to rather actively learn about the underlying feature weights than about the feature rankings alone. Our results constrain the types of decision strategies people might apply, although they do not limit them to just the two edge cases of full information or heuristic strategies. Instead, a more subtle picture emerges, where even if participants’ choices corresponded to a TTB heuristic during decision-making at test, their active learning behavior can still be governed by a feature-based algorithm during self-directed learning.

In summary, the research presented here makes the following main contributions:

1. Our primary intend is a methodological proposal: We put forward model-based active learning as a general-purpose model comparison technique for cognitive science, and demonstrate its usefulness in a decision making example.
2. We derive model-based active learning versions for both Take-The-Best and logistic regression and compare these in an active learning experiment, assessing whether participants’ self-directed learning is more in line with an active version of TTB or logistic regression.
3. Our experimental findings suggest that participants’ active learning behavior is better described by a model-based version of logistic regression than TTB, which suggests people focus more on learning feature weights rather than feature ranks.

Our findings also make a theoretical advance, demonstrating that people can sometimes learn more complex representations than what is visible in more traditional tests of overt behavior.

## Traditional model tests of heuristics

While the fast-and-frugal heuristics approach gained popularity based on showing statistical less-is-more effects when producing out-of-sample predictions for multiple real world data sets (Brighton, 2006; Chater, Oaksford, Nakisa, & Redington, 2003; Czerlinski et al., 1999), the empirical evidence for any specific use of heuristics is still controversial. One of the core questions in this debate concerns the way in which people look up and integrate information, and whether this behavior adheres to the search and stopping rules imposed by heuristics or full-information strategies (Gigerenzer, Todd, ABC Research Group, et al., 1999; Newell & Shanks, 2003). The main assumption behind this approach is that if people were using TTB, their information search of features should stop in accordance with TTB’s stopping rule. In contrast, if they were using a full-information strategy, information search should continue beyond that. Interestingly, a large body of studies produced results contradicting TTB’s predictions - participants tend to look up more information than what is needed when utilizing TTB (Bröder, 2000; Glöckner & Betsch, 2008; Newell & Shanks, 2003; Newell et al., 2003). Other studies find that people’s search behavior and response times conform to the TTB heuristic at least sometimes, and in an adaptive manner (e.g. Bergert & Nosofsky, 2007; Bröder & Gaissmaier, 2007; Dieckmann & Rieskamp, 2007; Rieskamp & Dieckmann, 2012).

However, using people’s additional feature integration as evidence for either cognitive model has limited explanatory power. That is because presenting people with given features at hand, e.g., on a screen in front of them, has been argued to elicit different cognitive processes than the natural heuristic decision making process, which is assumed to rely on learned feature knowledge from memory (sometimes called *inference from memory* versus *inference from givens*, see Gigerenzer & Goldstein, 2011). Secondly, additional feature search is only a limited approach to the problem at hand since it is only ever possible to check for *k* + 1 feature look-ups given that *k* features have been presented so far, and it has been argued that people might even look up additional information without utilizing it later on (Marewski & Mehlhorn, 2011). Other studies tried to elicit people’s use of the TTB heuristic by introducing additional information search cost to acquiring feature information. Dieckmann and Rieskamp (2007) showed that stopping search in accordance with TTB was more than twice as prevalent when the features’ look-up cost was increased, where the prevalence rate also depended on the redundancy among features. Yet, how much does this finding tell us about people’s actual use of TTB? Although the finding may be partially explained by an adaptive use of TTB, it is also sensible to use less information when features are costly, i.e., people did what a resource-rational account of decision making would also predict (cf Lieder, Krueger, & Griffiths, 2017). Much stronger evidence for the use of frugal TTB strategies would come from people readily using TTB without introducing additional search cost, i.e., a preference for heuristics even if information is free.

By far the most common method of pitting full-information models and heuristics against each other have been statistical simulations. Heuristics have been shown to outperform full-information models when making predictions within diverse data sets (Czerlinski et al., 1999; Gigerenzer & Goldstein, 1996; Katsikopoulos, Schooler, & Hertwig, 2010). Other studies have shown that there is no strong reason to prefer TTB over other cognitive models (Chater et al., 2003). However, just because one model class generates better out-of-sample predictions, it does not follow that it is also a descriptive model of human behavior. Beyond that, when one’s goal is to achieve high predictive accuracy at capturing people’s decisions, there is no strong evidence for preferring frugal heuristics over other strategies (Schulz, Speekenbrink, & Shanks, 2014). Indeed, *actually* learning a non-compensatory strategy using only heuristic building blocks turns out to be a non-trivial computational challenge (Schulz, Speekenbrink, & Meder, 2016). Nonetheless, evidence from statistical less-is-more effects is frequently used as evidence for psychological less-is-more effects, as evident in the literature: *“More information or computation can decrease accuracy; therefore, minds rely on simple heuristics in order to be more accurate than strategies that use more information and time*.” (Gigerenzer & Brighton, 2009, p.110).

In sum, existing model testing procedures are lacking in external validity, psychological plausibility and discrimination accuracy. For example, binary forced choice tasks are not good at discriminating *how* people arrive at their choice, i.e., by weighing and adding features, or by relying on the feature rank orders and the highest ranked cue (also in part because heuristics and regression models often make the same predictions). Instead, we believe that psychological processing needs to be investigated *in situ* and with appropriate methods that can elicit people’s representations concurrently.

We propose active information sampling as a sensitive method to assess which cognitive model people use. Our method is useful insofar as it focuses on how the information is learned in the first place, and how it is subsequently used in decision making. Using self-directed learning as a window onto the implementation of cognitive strategies, it becomes possible to set up active learning algorithms for many models of decision making, an approach we refer to as *model-based active learning*.

## Model-based active learning

The main idea behind psychological theories of active learning is to describe a learning agent as efficiently designing experiments (Chaloner & Larntz, 1989; Coenen, Rehder, & Gureckis, 2015; Gureckis & Markant, 2012). This means that, given that she wishes to find the true hypothesis out of many potential explanations as fast as possible, an agent assigns prior probabilities to each hypothesis according to some objective criterion such as available frequency data or according to the subjectively judged plausibility of each hypothesis. Each possible outcome of each possible experiment can then be considered in a “preposterior analysis” (Schlaifer & Raiffa, 1961), assessing the ways in which each possible experimental outcome could modify beliefs about the hypotheses. Optimal experimental design relies on maximizing an informational utility, which is typically a measure of how much the beliefs about the hypotheses have changed and how much that change has reduced uncertainty. A common measure of uncertainty reduction is the reduction of entropy (Shannon, 1948). Entropy expresses the prior uncertainty about a hypothesis, while the reduction in entropy refers to the reduction in uncertainty about the hypothesis after seeing some evidence (Nelson, 2005).

There has been a great deal of interest in both normative and descriptive questions surrounding human information acquisition (Nelson, McKenzie, Cottrell, & Sejnowski, 2010). In a probabilistic framework, active learning approaches have been used to model human behavior on cognitive tasks such as feature learning (Griffiths & Austerweil, 2009), reward-specific information search (Meder & Nelson, 2012), and to assess the trade-off between exploration and exploitation (Knox, Otto, Stone, & Love, 2011; Schulz, Konstantinidis, & Speekenbrink, 2017). Oaksford and Chater (1994) were among the first to define participants’ information selection behavior as active information acquisition. In a series of experiments they showed that the way people select cards in the Wason card selection task (see Wason, 1966) is in line with predictions derived form optimal experimental design principles, thereby redefining what was previously thought of as irrational behavior as a sensible strategy to test hypotheses. Markant and Gureckis (2014) found that it is more efficient to select rather than to receive information when testing hypotheses about categories and that participants tend to search in high information regions along the category boundaries when actively selecting information. This in turn led to faster learning and lower classification error (Markant & Gureckis, 2014). Lagnado and Sloman (2004) found that participants learn about causal structure more easily if they are allowed to perform timely interventions as compared to just passively observing the system. This idea that was later expanded by Bramley, Lagnado, and Speekenbrink (2015) and Coenen et al. (2015) to account for participants’ active causal learning behavior.

The goal of the current paper is to understand to what extent uncertainty-reducing active learning implementations of classic decision strategies can describe participants’ active information search behavior, in order to distinguish among two prominent decision making strategies, namely Take The Best and a weighted additive strategy (logistic regression). In order to do so, we use the notion of efficient information gathering to assess what people are trying to learn when performing self-directed sampling (as also suggested by Bramley, Dayan, Griffiths, & Lagnado, 2017). As active learning counterparts to these decision making strategies have not yet been proposed, we develop two learning algorithms which aim to maximize information gain, one for a feature-ranking and one for a feature-weighing strategy. While we focus on these two strategies to address whether people learn to develop heuristics or full-information strategies, it is important to note that our general model-based active learning framework can incorporate many other classes of psychological models.

### Active learning algorithms

Both active learning algorithms essentially rely on a one-step ahead greedy approach of minimizing, over all possible input queries, the expected posterior uncertainty about a set of competing hypotheses. Input queries are all those test stimuli that an active agent chooses from to learn about the underlying hypotheses, e.g., turning a card in the Wason selection task, or asking a question about a category object such as a cat that resembles a dog. In our experiment, a query refers to a binary comparison between two Alien species to find out which one wins in a competition. Greedy algorithms always choose as the next pair the query which promises to maximally reduce the uncertainty immediately following feedback on the chosen input query (i.e., focusing only on the knowledge gained from a competition between the chosen pair of aliens, but not on what could be learned thereafter).

#### Active Take-The-Best

The TTB heuristic assumes that people look up features sequentially in the order of their validity, and stop this search as soon as a feature discriminates (favors one option over the other). Therefore, the active TTB algorithm learns with the goal of establishing the correct feature validity rank order, rather than the precise validity weights. We implement a Bayesian version of TTB that estimates a distribution over feature weights via Metropolis-Hastings sampling, thereby generating multiple realizations of the TTB heuristic given its current posteriors’ feature rank orders. In particular, using the data of pairwise Alien comparisons seen so far, this algorithms fits a Bayesian linear regression where the comparison outcomes are regressed onto each feature individually, assuming independent features as in heuristic feature validities (Martignon & Hoffrage, 1999). Afterwards, we sample realizations from each of the distributions and treat them as a proposed feature rank order for one proposed TTB model. This process is repeated multiple times and we call the multiple realizations of TTB *proposal TTBs*. These proposals can differ in their feature rank orders as they are based on Monte Carlo samples. The proposal TTBs are then used to create multiple realizations of model predictions over the input queries (i.e., possible pairwise comparisons), and the predictive variances for input queries can be assessed as the amount of disagreement over all proposal TTBs. Higher disagreement means that the different proposal TTBs generated a higher predictive variance for an input query. Queries with higher uncertainty lead to higher disagreement and are therefore (in the long run) expected to reduce uncertainty more strongly. The active TTB algorithm therefore chooses that query as its next observation where the uncertainty is currently highest, as this is the query where uncertainty reduction is expected to be large. Thus, the active TTB algorithm makes choices that are expected to learn as much about the underlying feature rank orders as possible and therefore values queries more whose rank-based results are currently uncertain.

#### Active logistic regression

Logistic regression is set up as the competing full-information model. In contrast to the heuristic which relies on the feature rank order and ignores the feature weights, logistic regression weighs each feature and integrates all of them. The active logistic regression algorithm was set up to learn with the goal of establishing the feature weight magnitudes. We implement a Bayesian version of logistic regression based on a random walk Metropolis algorithm. We use Gibbs sampling to generate posterior MCMC-samples over the regression weights. In particular, we perform Bayesian logistic regression that takes *all of the features* as the independent variables and the outcomes of the comparisons as the dependent variable. This leads to a posterior distribution over different weights within the same regression model. Afterwards, we sample realizations from the posterior distributions and treat them as the proposed weights for one proposed logistic regression model. This process is repeated multiple times and we will call the multiple realizations of logistic regression *proposal logistic regression models*.

The proposal logistic regression models can then be used to generate model predictions over all input queries by sampling realizations for the weights. As in the active TTB model, the predictive variance for each query can finally be approximated by the disagreement among proposal models. Thereby we build a logistic regression analogue to the active TTB that is conceptually similar, i.e. it queries inputs whose output is currently uncertain. The active logistic model hence chooses that input query next which has the largest predictive variance and is expected to reduce uncertainty as much as possible. Instead of trying to drive down the uncertainty with respect to feature rank orders, the logistic algorithm tries to drive down uncertainty with respect to the regression weights.

## Creating environments of different compensatoriness

We are interested in the performance of the two proposed active learning models in environments with different “compensatoriness” (Martignon & Hoffrage, 1999). Decision strategies are assumed to perform best in matching environments that have the same properties, i.e., compensatory strategies performs best in a compensatory environment and TTB in a non-compensatory environment (Martignon & Hoffrage, 1999). Note that a non-compensatory environment can be defined as a logistic regression environment in which the *β* weights are exponentially decreasing. In order to create different degrees of “compensatoriness”, we make use of a mathematical trick that allows us to rely on a single parameter to smoothly vary from compensatory to non-compensatory environments through a “stick breaking process”.

If we have 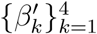, then we can take

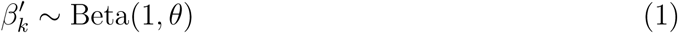

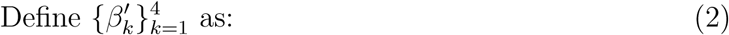

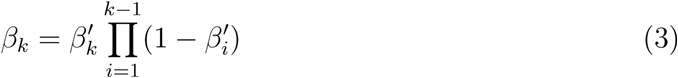

to produce differently sized breaks from a stick of unit size. As the expectation of the Beta-distribution is defined as 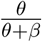, we can infer that an environment where Take-The-Best would thrive can be created by setting *θ* to 1 as this would lead to an expected stick break of half of the stick per break, i.e. very non-compensatory. As *θ* gets bigger, the result are more and more uniformly distributed feature importances. The bound that separates compensatory from non-compensatory strategies is *θ* = 1, and we use *θ* = [0, 0.5,1, 2, ∞] for all the upcoming scenarios to cover a range of compensatory/non-compensatory environments.

The heuristic literature predicts people’s choice of decision model is adaptive to compensatoriness and we sought to see whether this is also the case for active learning models Martignon and Hoffrage (1999).

## Experiment

We designed an experiment to find out whether people are more likely to follow a rank-based or a weight-based active learning strategy. We hypothesized that the active learning model that best describes people’s information queries would also constitute their most likely cognitive decision strategy. We also wanted to investigate whether people are sensitive to the structure of the environment (the degree of compensatoriness) in their active queries, such that in non-compensatory environments participants would be better matched by active TTB, while in compensatory environments they would be better matched by active logistic regression. Therefore, we assigned people randomly to one of the five above-mentioned compensatoriness conditions.

### Participants

Participants *(N =* 264, 137 females, average age *M =* 35.4) were recruited via Amazon Mechanical Turk to take part in an “Alien Olympics” study. Participants were paid $0.50 for participation plus an additional bonus between $0 and $0.50 depending on their performance.

### Procedure and stimuli

The experiment was divided into a learning and test phase. The learning phase consisted of participants actively choosing Alien pairs to fight against each other, with as binary outcome one of the aliens winning, and the other losing the match (i.e., a draw was not possible).

Aliens varied on four different features (see Figure 2), resulting in a total of 16 possible Alien types in the Alien database for the entire experiment. Participants were told that the presence of each feature was helpful in a fight, e.g., wings enabled an Alien to fly which helps in attacking enemies, while camouflage is useful for hiding from enemies, and antennae provide surrounding vision. The helpfulness of features was explained to participants at the start of the experiment and they were also told that the different features might not all be of equal importance for an Alien’s strength in a fight.

**Figure 1.**
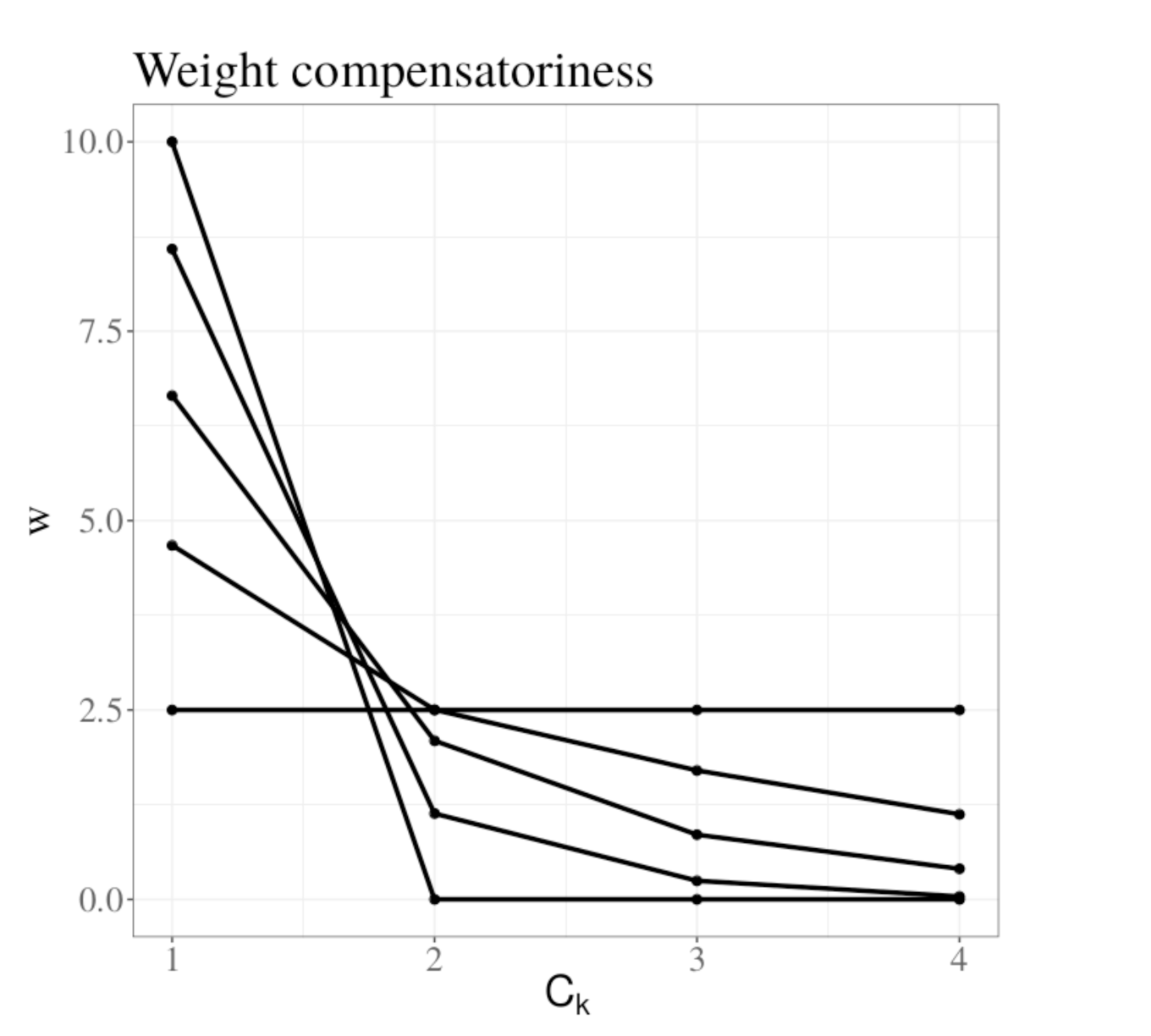
Compensatoriness for five different levels of *θ*. The x-axis represents four different features and the y-axis displays the feature weight magnitudes. These five levels of compensatoriness were used as five conditions in the Experiment below.

**Figure 2.**
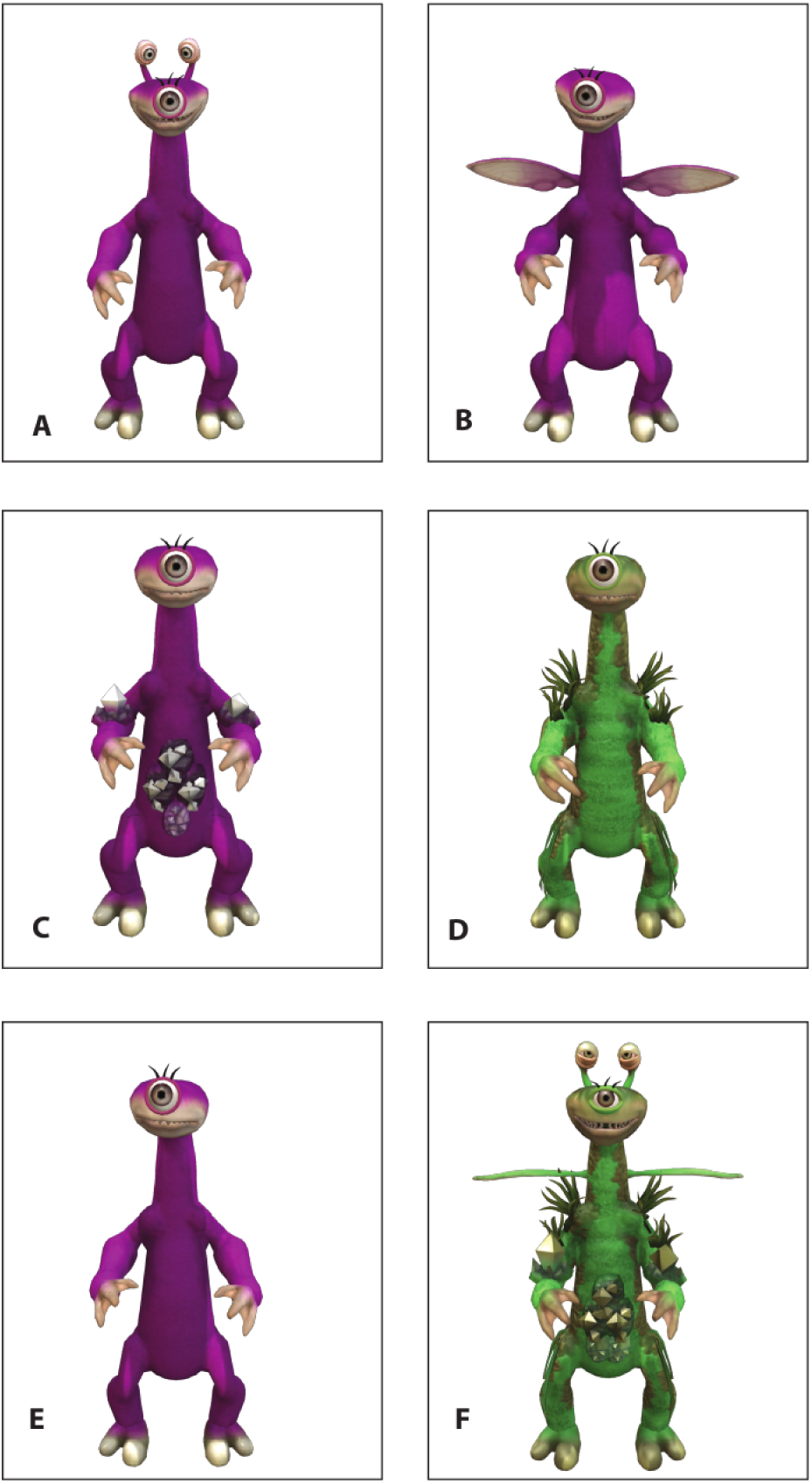
Aliens varied on 4 different features (A-D): Antennae, Wings, Diamonds, and Camouflage. E: Alien without features, F: Alien with all features.

#### Learning Phase

On each learning trial, participants were presented with four Aliens on the screen and had to choose which pair of Aliens would fight against each other on that given trail.

We emphasized that participants should pick Aliens wisely by selecting informative comparisons, as the goal was to learn how the different features influenced an Alien’s chances to win. Participants were informed that they would need this feature knowledge later on for the assessment task (the test phase). The presented four Aliens were randomly sampled from the 16 Alien types without replacement. With four Aliens on any given learning trial, there are six possible pairwise comparisons to choose from. After selecting a pair, participants received feedback about which Alien had won the competition. Reflecting the probabilistic nature of the outcomes, participants were also told that, as in any sport, sometimes a weaker Alien could win against a stronger competitor. The underlying feature weights that determined the strength of the aliens depended on the compensatoriness condition a participant was in (as shown in Fig. 1). The actual outcomes observed in feedback were generated by using the weights (standardized to always add up to 10) and applying logistic regression in order to determine the difference in strength between a pair of Aliens (i.e., the likelihood of one winning against the other Alien). The learning phase consisted of 30 trials.

#### Test Phase

The test phase was designed to assess what people had learned during the learning phase and was structured as follows: On each test trial, participants were presented with two different Aliens that were again randomly drawn from the Alien database without replacement.

We told participants that these Aliens were the candidates for their “Olympic Team”, and it was their task to always choose the Alien they considered to be stronger based on what they had learned about the features in the learning phase. The test phase consisted of 10 trials forcing participants to make binary choices. Participants were reminded that a bonus payment would depend on their performance in this test phase. The bonus was calculated based on the average probability of their chosen Aliens in the test set to win against the other presented Alien. If 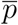 indicates that average probability over all test trials, then participants’ overall reward was calculated as 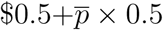.

### Results

We present the results of the test phase first before moving onto the active learning results. We will refer to the results from the test phase as “passive” model fits and to the learning phase as “active” model comparison.

The average percentage of correct choices made was 74% with a range of [30%, 97%]. Participants’ performance at identifying the stronger Aliens during the test stage was highly above chance, *t*(263) = 27.44, *p <* 0.001. Performance varied as a function of the compensatoriness condition that participants were in. Figure 3 represents the average performance score at test as a function of compensatoriness: As the environmental structure becomes more non-compensatory (i.e., more weight on just a few features), the average performance drops. This is intuitive as there is less information to be learned when one feature dominates all others, a scenario which causes similar probabilities of wining across Aliens and therefore makes informative comparisons less frequent. However, peak performance was not observed for a fully compensatory environment, but rather for slightly less compensatory situations (*θ* = 2).

**Figure 3.**
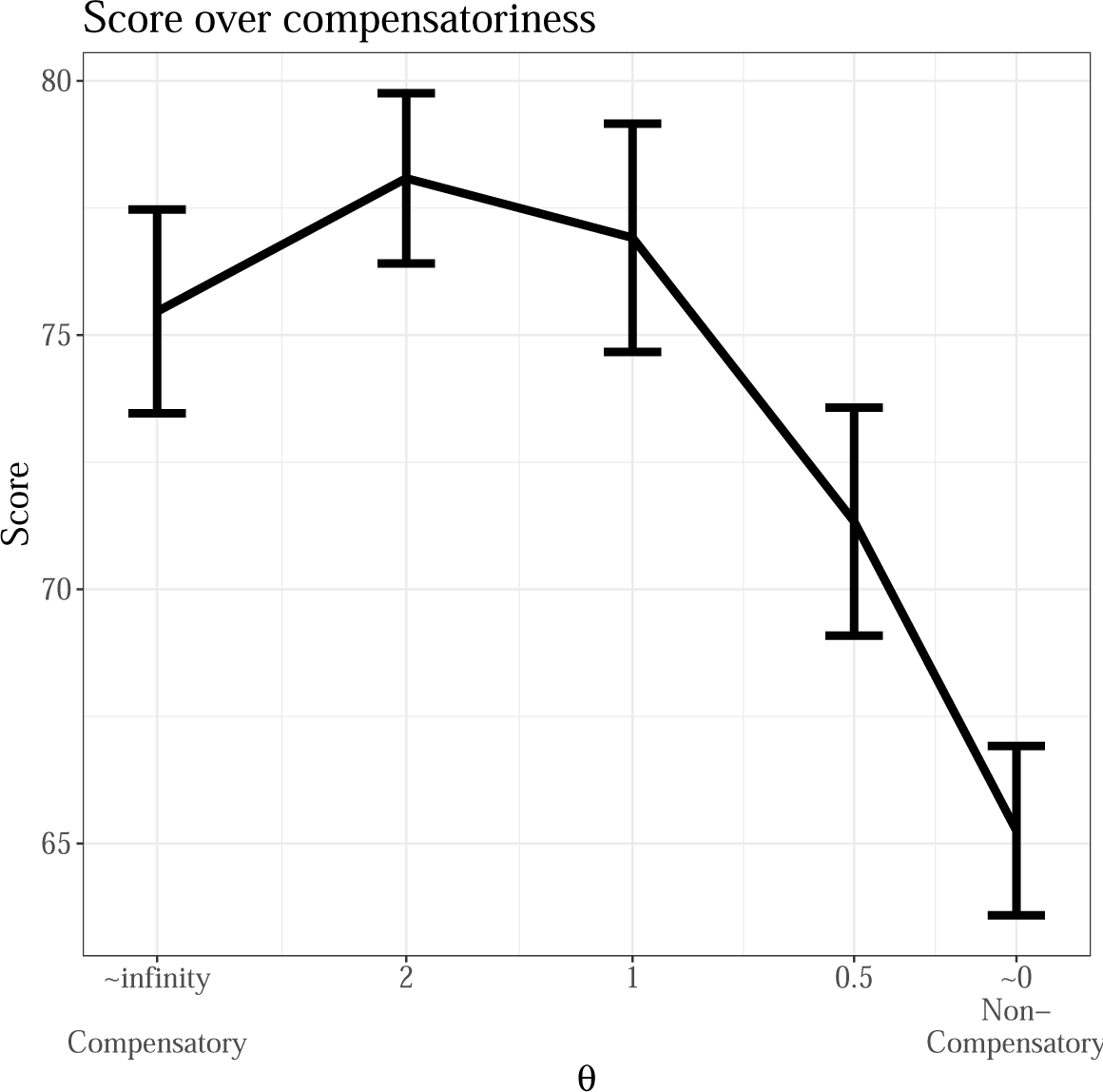
Average test performance by compensatoriness conditions. Y-axis represents the percent of correct choices that participants made across the 10 test trials. Error bars represent the standard error of the mean.

#### Passive model comparison

Next, we compare the performance of both the TTB heuristic and logistic regression at capturing participants’ behavior at test. The models used here to predict test choices represent the standard TTB heuristic and logistic regression model as found in the psychological literature (e.g., Czerlinski et al., 1999; Gigerenzer & Brighton, 2009), and are not to be confused with the active learning versions specified above.

We use a cross-validation procedure in which both algorithms learn weights from the Alien pairs in the learning phase (training sample) and make binary predictions for the Alien pairs in the test phase (test sample). A training sample consists of the set of pairwise alien comparisons a participant chose during learning and its respective feedback, while the test sample consists of the test trials the participant saw. Consequently, each participant has a different training experience depending on what queries they selected during the learning trials (and the attached feedback these selections produced). Both the TTB and logistic regression model are fit to each participant’s training sample, and the fitted weights are used to make predictions for participants’ test samples.

Figure 4 presents the predictive accuracy of the TTB and logistic regression model in comparison with a model that predicts at random. It can be seen that both logistic regression (*t*(243) = 51.43, *p <* 0.001, *d* = 3.17) and TTB (*t*(243) = 9.34, *p <* 0.001, *d* = 0.57) were better than the random model at predicting people’s behavior at test. However, logistic regression performs better than the TTB heuristic at capturing participants’ choices (*t*(243) = 15.83, *p <* 0.001, *d* = 0.97). These results suggest that participants’ decisions at test were more in line with a logistic regression strategy. If we were to stop here, the conclusion would be that the full-information model is psychologically more plausible than the heuristic. In the literature, most studies do not go beyond this point and rely on behavioral model fits assessed in the test phase (i.e., as indicated by percentage correct, R-squared, AIC, and so forth) to draw conclusions about people’s decision making processes. However, we argue that this approach represents a somewhat impoverished view on people’s inference processes. This common method to perform model comparisons predicts participants’ choices in a highly controlled and passive environment, where stimuli have been pre-selected by the experimenter and people do not get any freedom in choosing the stimuli that make up the binary choices.

**Figure 4.**
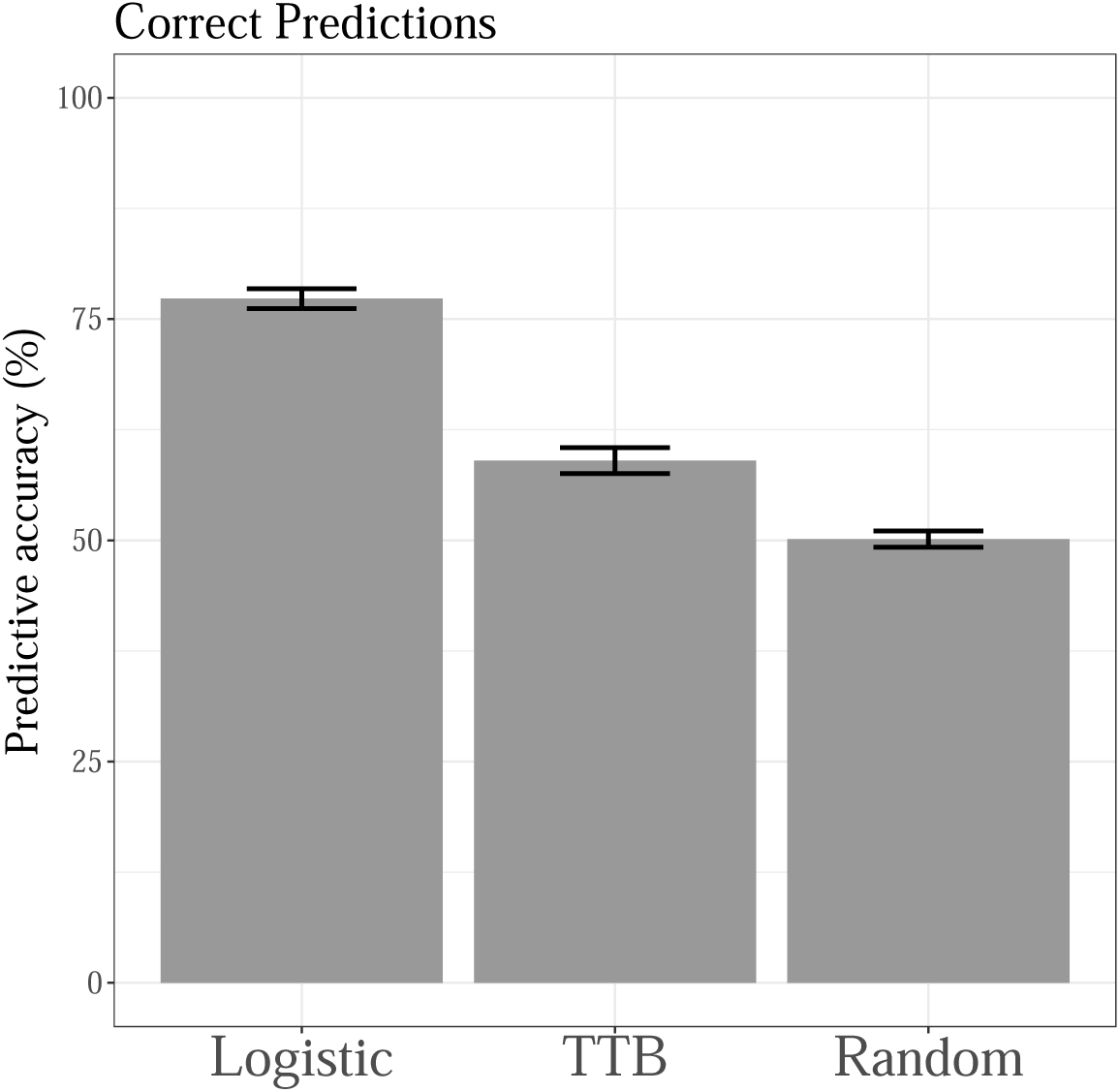
Predictive accuracy of the TTB heuristic, logistic regression, and a random model at test. Y-axis represents the percent of correct predictions with respect to peoples’ choices across the 10 test trials. Results are averaged across compensatoriness conditions. Error bars represent ± SEM.

#### Active learning queries

We focus on people’s active learning queries; we categorized all possible queries into the 8 subtypes that can be seen on the x-axis of Figure 5. For example, +000 signifies a comparison of two Aliens with three equal features (0 for draw), where one Alien had one more feature than the other Alien (+ for advantage). A +− 00 query compares two Aliens that are matched on two features but differ on two other features (+ for advantage, − for disadvantage). An example of +− 00 may be where one of the Aliens features “Wings” while the comparison Alien features “Camouflage”, but both possess “Diamonds” and “Antennae”. This query therefore indicates an assessment of whether “Wings” or “Camouflage” were more important for the outcome in an Alien fight. In general, it can be said that all queries, i.e., even queries such as + + +0, exhibit some information about feature weights, while for a rank based strategy (TTB) the best queries to learn about feature order are primarily the queries +− 00, + +−0, and + + +−.

**Figure 5.**
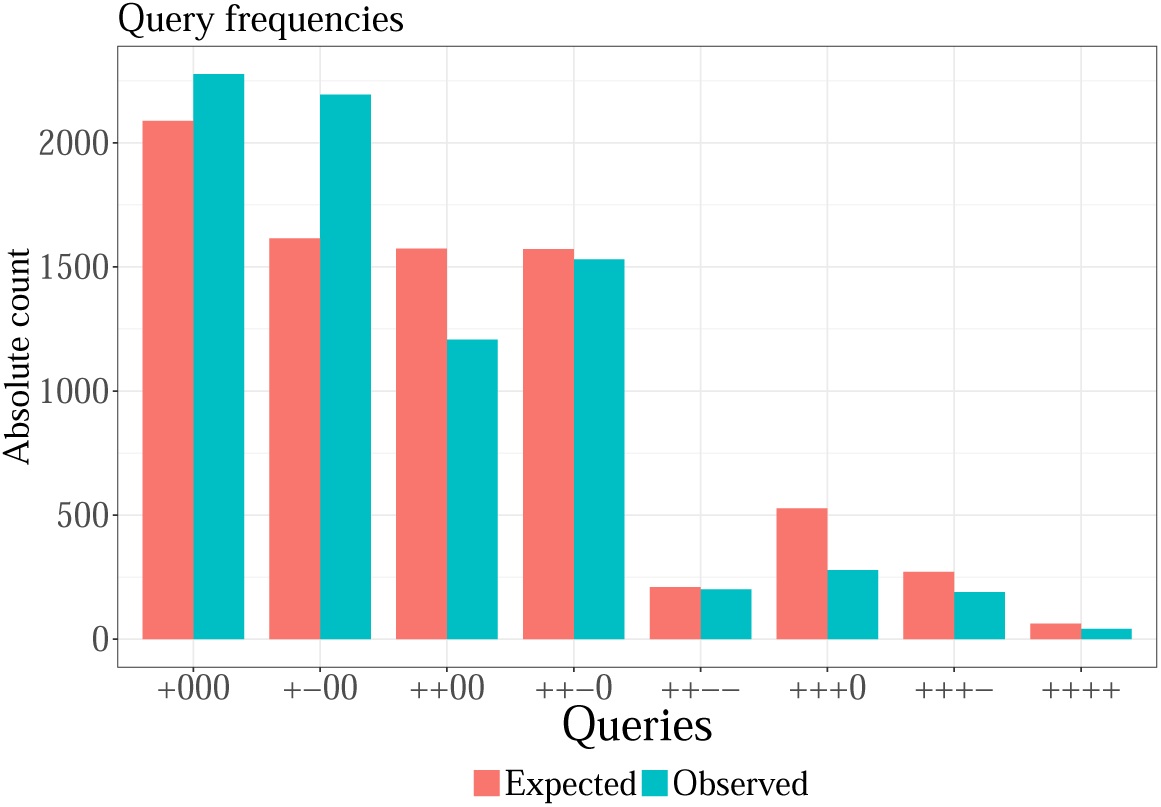
The 8 subtypes of active learning queries that participants could make. The y-axis represents the frequency of choosing the query across the 30 learning trials across 264 participants. Coding is as follows: ‘+’ = Alien has a feature that the other alien does not have (advantage); ‘-’ = Alien lacks a feature that the other alien has (disadvantage); ‘0’ = Both aliens have the feature, or both aliens do not have the feature (draw). The figure plots the observed frequency of choosing a particular query against the expected frequency of choosing a particular query under the base rate (probability of occurring naturally) in the experiment.

It can be seen in Figure 5 that people rarely chose comparisons where it is unclear what feature was responsible for an outcome, e.g., a comparison of an Alien with 3 or 4 more features than its competitor (i.e., + + +0 or + +). Instead, the most common comparisons was a more controlled comparison, such as the +− 00. Crucially, to interpret people’s queries in a meaningful way, one needs to look at how people’s queries differ from what would be expected if they chose the pairs randomly, i.e., under the experiments’ base rate. Every query has a different base rate probability of occurring on any given learning trial, i.e., some Alien comparisons are more likely than others due to the nature of how the 16 different Alien types were generated. Thus, we compared the absolute frequencies with which participants chose each query across the learning phase (“Observed” in Figure 5) with the probabilities of each query occurring at any trial for any participant. The base rate probabilities were measured as the relative frequency of query occurrence across the full experiment. A Chi-square test of goodness of fit reveals that the observed distribution of frequencies was significantly different from the expected frequency distribution under the base rate probabilities, *χ*^2^(2) = 460.23, *p* < 2.2 × 10^−16^, *ϕ* = 0.23. Hence, it can be concluded that people’s behavior in the learning task was non-random.

Comparing the observed frequency counts against the expected frequency counts under the null hypothesis of random choice (Figure 5) reveals that the biggest difference was that participants queried the +000 and the +− 00 comparisons a lot more, while the + + 00 and the + + +0 a lot less, then expected under random choice. This suggests that participants were choosing more informative comparisons than expected, where the source of an Alien’s strength can be clearly attributed to a feature. Although participants were told in the instructions that all features are helpful in fights, the +000 query was still a relatively popular query, which essentially tests whether a feature has a positive or negative effect on an Alien’s strength -a sensible query if the features valences were assumed to be unknown. Moreover, repeating this query multiple times leads to an increased knowledge about the magnitude of a feature’s overall weight (i.e., its validity).

The query that shows the largest difference from the base rate is the + − 00 query. This test involves comparing two Aliens which differ on 2 features, for example “Wings” and “Camouflage”, and assessing the relative importance of Wings in comparison to Camouflage for the outcome directly. As mentioned above, this query can be seen as a test for establishing feature order, but performing this query multiple times can also assist in learning differential weights.

Finally, we assessed changes in query over time, to see if people may apply different strategies at different stages in the learning phase. Figure 6 plots query frequencies as a function of learning trial. What is apparent from this Figure is the general increase in in the +000 query, and a relative decrease in the other queries, including the + − 00 queries. At the final trial, the +000 query is the most common one, which goes against the base rate which would predict the + − 00 query to be less frequent than the +000 query (Figure 5).

**Figure 6.**
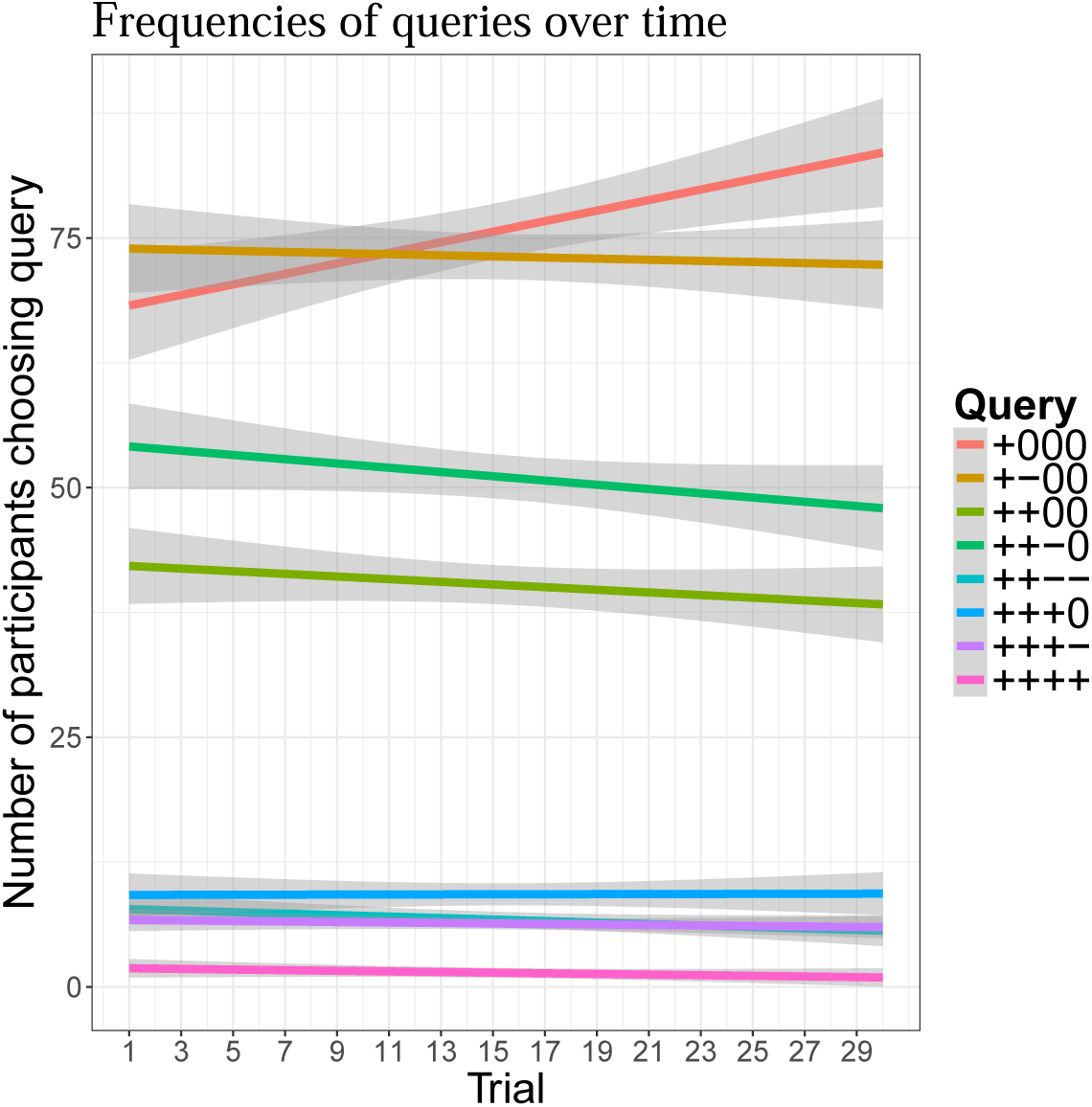
The 8 subtypes of active learning queries that participants made as a function of time, i.e., learning trials progressing from 1 to 30 (x-axis). The y-axis represents the number of participants out of 264 participants in total that chose the query at each trial. The data points for each graph were smoothed to a line graph with a generalized additive model as the smoothing function, where the boundaries represent the 95% confidence band.

It appears that people are systematically choosing + − 00 more often up to a point, while later in the experiment, e.g., approximately from trial 21 onwards, the simpler +000 query starts to dominate (the slope for the +000 query is significant with *β* = 0.53 ± 0.16, *p <* .01). Elaborating on the meaning of the +000 query, it seems to be a relatively uninformative query at first sight, especially once the features’ valences are established by the learner. However, repeating this query multiple times will reveal a feature’s overall relative importance and therefore can indicate a further focus on learning the weight of this feature. Therefore, the increase in the +000 query in later trials may reflect a preference to perform more focused tests towards the end. This is in line with recent research indicating people have a tendency to to build up models gradually over trials, with more fine-grained testing towards later learning trials (Bramley et al., 2017). Other research also suggests people’s active learning strategies may often use a combination of discriminatory and confirmatory testing, e.g., in causal intervention studies (Coenen et al., 2015). Lastly, it can be said about Figure 6 that participants overall preferred the more complex + + −0 query to the + + 00 query (middle green lines), despite equal base rates. The + + −0 query can be seen as a more complex test, comparing an Alien with two features to an Alien with one other feature, e.g., assessing how much better or worse “Antenna” is compared to “Wings” and “Diamonds” combined.

#### Model-based active learning comparison

Finally, we look at the correspondence between the active learning algorithms and people’s queries. We let both the TTB and logistic regression algorithms learn in the same compensatory and non-compensatory environment as the participants, by creating as many simulated participant profiles as there were participants in each compensatoriness condition. Then, we let the models learn over time. In particular, the models are fed participants’ data at time point *t* and generate one-step ahead predictions for trial *t* + 1. Both active algorithms are driven by uncertainty sampling, which means they are predicted to choose that Alien comparison next which maximally reduces uncertainty. If, for example, there are six possible Alien comparisons to choose from at a given learning trial *t* + 1, the algorithms produce a vector of uncertainties for all possible queries at time *t* + 1, and the query producing the greatest uncertainty would be chosen. To assess correspondence between participants’ choices and the algorithms, participants’ actual choices across all learning trails were regressed onto the predicted model-based uncertainties — normalized per trial — in a logistic regression for each participant individually.

Figure 7 shows the match between the active algorithms and participants’ queries, as measured by the resulting averaged McFadden’s pseudo R-squared (McFadden et al., 1973), as a function of compensatoriness conditions. Results demonstrate that the active logistic algorithm captured people’s queries better than the TTB algorithm across all environments. This was true regardless of the compensatoriness condition. These findings are in line with our predictions and the results from test phase in Figure 4. We ruled out that people applied a confirmatory sampling strategy by explicitly testing an active learning model that chooses queries which minimize uncertainty reduction, which does no apply here as can be seen from the non-negative correlations in Figure 7.

**Figure 7.**
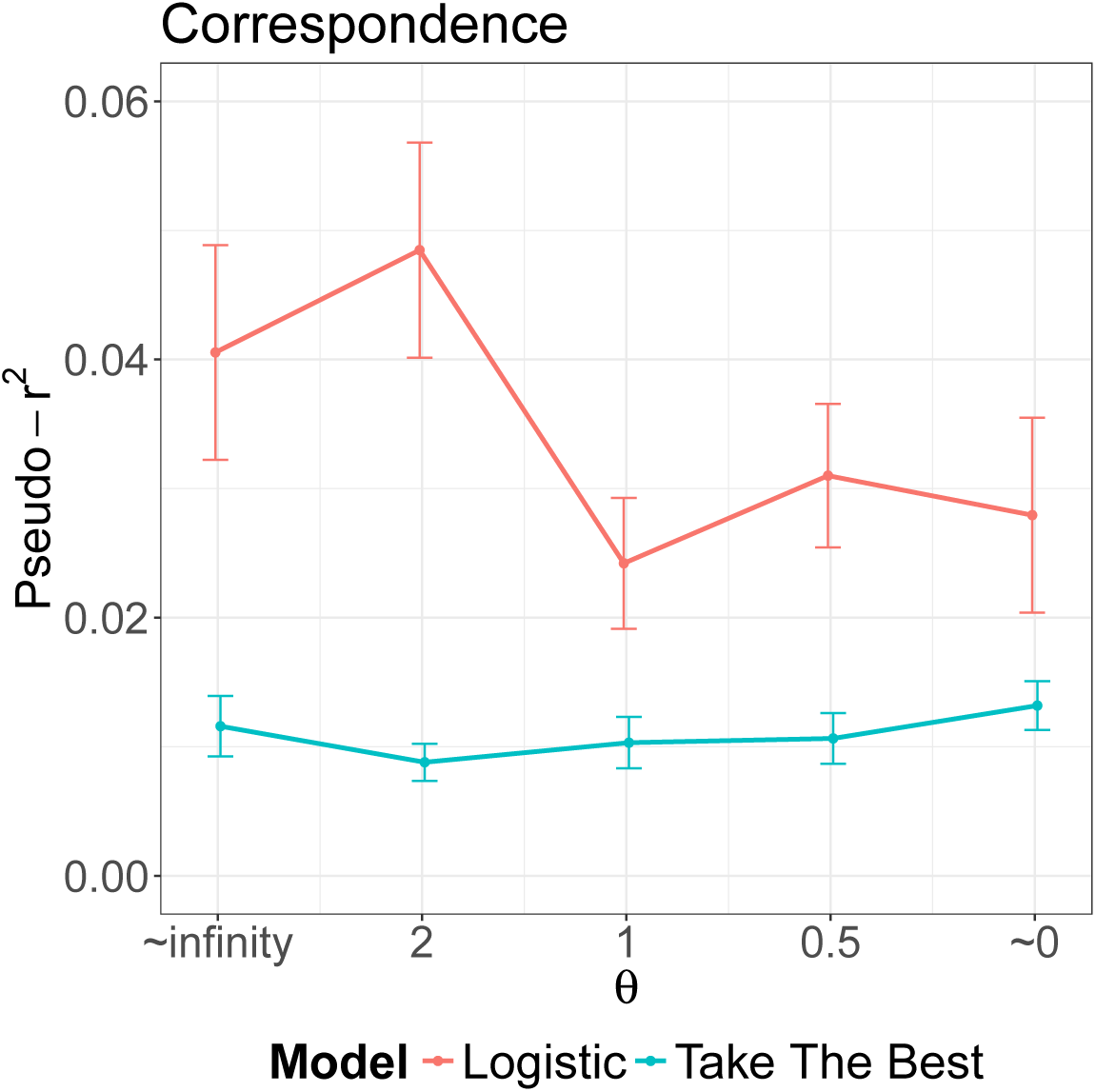
Correspondence between the active learning algorithms and participants’ active queries, as a function of the compensatoriness condition. Results are established from simulating active participants with both active learning models in a step-by-step fashion predicting participants’ next queries. The active logistic algorithm was better at capturing people’s active queries than the TTB algorithm, regardless of compensatoriness condition. Error bars represent ± SEM.

While the pseudo R-Squared values may appear to be low relative to traditional passive model fits, two things need to be noted. Non-active behavioral model fits typically predict one out of two objects to choose from (e.g., one of two Aliens in a test pair), while there are typically many active queries to choose from (e.g., six possible queries on each trial in our experiment), and these can be of roughly equal informativeness (uncertainty) to an ideal learner, meaning they could be tested in any order. Hence this automatically reduces predictability of active queries compared to standard decision making tasks and has been reported in other studies (e.g., see Bramley et al. (2017) for further discussion). Secondly, we did not optimize hyper-parameters to enforce a greater model fit of the model-based active learning algorithms as is frequently done in other active learning studies. Instead, all predictions were purely derived from what a (close to) rational active learner would value on every trial, had she learned with one particular decision strategy in mind and seen a participant’s queries up to that trial. Thus, although these measures are small, our results nonetheless indicate that participants took the uncertainty of the different queries into account, thereby significantly valuing uncertainty reduction over confirmatory hypothesis testing.

Interestingly, no clear relationship between the compensatoriness among features and the best fit active models can be seen, e.g., it was not the case that the active TTB model better predicted people’s queries in non-compensatory conditions and the active logistic model better fit in compensatory conditions. Instead, people seem to be learning the weights regardless of compensatoriness. These results go against parts of the heuristic literature claiming people are not able to learn weights due to capacity limitations and instead rely on heuristics which are less computational demanding (Gigerenzer et al., 1999). Particularly, in the highly non-compensatory environments which are ideal for the TTB heuristic (Martignon & Hoffrage, 1999; Rieskamp & Dieckmann, 2012), it may be surprising that people still behaved more in line with an active logistic regression strategy.

The aforementioned results are also confirmed by assessing the aggregated active model comparison results. In the active condition, the mean performance for both the logistic regression (AIC = 160.5, *t*(263) = 8.49, *p <* 0.001, *d* = 0.52) and the Take-The-Best model (AIC = 164.4, *t*(263) = 4.63, *p <* 0.001, *d* = 0.28) was better than a random model. However, the active logistic regression model described participants’ behavior better than the active Take-The-Best model (*t*(263) = 7.88, *p <* 0.001, *d* = 0.49). This suggests that participants were actively querying in a manner more consistent with a feature-weight based model.

**Table 1.** Number of participants best classified by an active TTB model versus active logistic model during the learning phase (columns), in relation to those best fit by a TTB model and a logistic regression model at test (rows). The classification is performed based on the lower AIC.

| | TTB active | LOG active | Σpassive |
| --- | --- | --- | --- |
| TTB passive | 47 | 67 | 114 |
| LOG passive | 43 | 107 | 150 |
| Σactive | 90 | 174 | 264 |

### Dissociation between active and passive learning

Table 1 shows a 2-by-2 frequency table displaying the number of participants best fit by either model (as indicated by lower AIC) for both the learning and test phase. The columns labeled “active” refer to the active learning algorithm applied to the learning phase, while the rows labeled “passive” refer to both the passive TTB and LOG model applied to the test phase, respectively. To the extent that there should be internal consistency between the learning and test phase, one would expect that the diagonal entries of this 2-by-2 matrix should be a lot higher than the off-diagonal entries. For example, a learner who is best described as a TTB user during decision-making at test would also be expected to be best described as a TTB learner during learning. Indeed, results show that the majority of people that were best fit by an active logistic model (LOG active), were also best fit by a passive logistic model (LOG passive) at test, i.e., 107 participants in the bottom right cell. This is good evidence for the internal consistency of an individual’s use of strategies in our experiment. However, while there was a clear majority of people better fit by an active logistic model compared to the active TTB model in the learning phase (Σactive: 174 vs. 90), the total number best fit by TTB passive and LOG passive at test was much more balanced (Σpassive: 114 vs. 150).

Most strikingly, some people that were best described by the LOG active algorithm during learning, appear to be best predicted by TTB passive at test, i.e., the 67 participants in the top right cell. This suggests that there were people that looked like they applied TTB at test, but when studying the active learning algorithms it becomes apparent that they were in fact also trying to actively learn the feature weights. This finding is particularly interesting as it may indicate that these participants empirically matched the TTB decision rule in making binary choices, but might have had the feature weights at their disposal. This result adds evidence to the fact that passive model comparisons are often not sufficient as a model discrimination technique and may even obscure psychological processing that could be uncovered with model-based active learning studies.

## General Discussion

Deciding between different options is common activity in our daily lives. Yet, definite answers to the question of whether people do so by following simple heuristics or differentially weighing and integrating the available information remain elusive. To overcome the existing conundrum of decision making strategies, we proposed to enrich the available repertoire of model comparison techniques by adding *model-based active learning*. Model-based active learning is built on the assumption that cognitive agents actively query information in the environment in order to minimize uncertainty about the cognitive model they utilize in that particular environment. This proposal is built on a rich canon of past research on human active learning (Bramley et al., 2015; Coenen et al., 2017; Markant & Gureckis, 2014; Nelson, 2005). Our work contributes to this growing literature in active learning by incorporating uncertainty sampling to determine which cognitive model best characterizes human decision making.

Using this approach, we built active learning versions of both the Take-The-Best heuristic as well as a logistic regression decision strategy. Whereas the former only aims to reduce uncertainty about feature rank orders, the latter tries to minimize uncertainty about weights. This critical difference allows model predictions to be disentangled in regards to information sampling. In our experiment, human participants selected comparisons to learn about the underlying task structure. Our results showed that for both active learning as well as the predictions at test, a logistic regression strategy described participants’ behavior better than the Take-The-Best heuristic. Thus, our newly-introduced method enabled us to find evidence for a preference of weight-based over rank-based decision strategies. These analyses expand the scope of model evaluation for theories of decision making.

The dissociation between people’s active learning strategies and their choice strategies at test (Table 1) suggests that people may be more sensitive to differential weighting of information than is evident from the more common procedure of passive model fitting to participants’ binary choices. This finding is in line with the possibility that heuristics may provide a good general characterization of data but that other accounts that are sensitive to additional information sources may perform even better (Parpart, Jones, & Love, 2017). Here, active learning provided a novel means to reveal the nuances of how people make decisions.

The current work is a first step towards a fully-fledged theory of model-based active learning. We hope that future research expands and improves upon our results. First, the produced pseudo R-squared values within our model comparison are relatively low, which we believe is due to not optimizing the hyper-parameters but solely relying on a model’s uncertainty to generate predictions. Whilst we chose to restrict the free parameters for simplicity, there is room for improvement and alternative models in future work. The present work demonstrates how learning models in which uncertainty is quantified can be evaluated in an active learning paradigm. Therefore, we welcome other researchers to evaluate other candidate models using the active learning blueprint we provide.

Secondly, the correspondence between the models used for active learning and the models used for the decision at test needs to be assessed further. Whereas our experiment has shown a relatively high consistency between the best active learning model and best model at test for each participant, there were also cases where this was not the case. This could either be the result of noise within the model comparison procedure or indicate that these participants might have used a different strategy for self-directed sampling than for decisions at test. For example, it is conceivable that participants start out learning about all the weights and —as soon as they learn that there is a non-compensatory structure— decide to ignore the weights and just base their decisions on more simplistic strategies (Niv et al., 2015). Further charting the landscape of correspondence between active and passive learning models will be another important step.

Our model-based active learning approach may also have interesting implications for the strategic sampling literature (Wilson, Geana, White, Ludvig, & Cohen, 2014). It is very difficult to assess whether an agent is an optimal sampler or not unless the analysis is done with respect to what model the learner is using. In line with this, our *model-based* learning approach puts particular emphasis on the model allowing for an optimal learner to actively learn in the first place, which a *model-free* uncertainty reduction approach cannot capture.

We believe that re-conceptualizing psychological models as active learning strategies will continue to provide insights to fundamental principles of learning and decision making, that would not be possible otherwise. We think this will apply to both the debate around heuristics, and hopefully other big debates in psychology. This, however, will require further replication and modifications of our current approach. Sometimes one should not make a decision after only one cue discriminates between two options.

## Acknowledgements

We thank Neil Bramley for helpful comments. PP and ES were supported by the UK PhD Centre in Financial Computing and Analytics at UCL Computer Science. This work is partly supported by the Leverhulme Trust grant RPG-2014-075 to BCL. ES is supported by a Postdoctoral Fellowship of the Harvard Data Science Initiative.

## Footnotes

1 If none of the features discriminate between the two options, then TTB picks one of them at random.

